# Structural instability of IкB kinase β promotes autophagic degradation through enhancement of Keap1 binding

**DOI:** 10.1101/407429

**Authors:** Mayu Kanamoto, Yoshihiro Tsuchiya, Yuki Nakao, Takafumi Suzuki, Hozumi Motohashi, Masayuki Yamamoto, Hideaki Kamata

**Author notes:** To whom correspondence should be address: Dr. Hideaki Kamata, Department of Molecular Medical Science, Graduate School of Biomedical Science, Hiroshima University, Kasumi 1-2-3, Minami-ku, Hiroshima, 734-8553, Japan.

## Abstract

IKKβ, an essential kinase of NF-кB signaling, is composed of an N-terminal kinase domain (KD) and a C-terminal scaffolding domain, containing a ubiquitin-like domain (ULD). The Hsp90 chaperon has special responsibility for folding of protein kinases including IKKβ. Here, we found that Hsp90 inhibition induced IKKβ degradation, which is partially mediated by Keap1. Geldanamycin (GA), a Hsp90 inhibitor, enhances association of IKKβ with Keap1 through the binding site in KD, and translocates IKKβ to detergent-insoluble fractions leading its autophagic degradation. An electrophile tBHQ suppressed Keap1-mediated proteasomal Nrf2 degradation but not autophagic IKKβ degradation. Substitution mutation of Leu353 to Ala in the ULD destabilizes IKKβ, enhances its association with Keap1, translocates it to detergent-insoluble fractions, and causes its autophagic degradation. These results suggest that Keap1 is involved in the degradation of structural destabilized IKKβ and negative regulation of NF-кB under proteotoxic stress.

## 1 Introduction

NF-кB is a critical transcription factor, regulating oxidative stress response, inflammation and carcinogenesis [1, 2]. In resting cells, NF-кB is inactivated by association with inhibitory proteins such as IкB*α* that prevent the DNA binding and nuclear transport. A vast array of stimuli including inflammatory cytokines and cellular stress activate NF-кB through the IкB kinase (IKK) complex, which consists of two catalytic subunits, IKK*α* and IKKβ, and a regulatory subunit NEMO/IKK*γ* [3-6]. While IKKβ mediates activation of the canonical NF-кB pathway in response to pro-inflammatory stimuli, IKK*α* has an indispensable role in non-canonical NF-кB signaling by phosphorylating NF-кB2 [7]. IKKβ is composed of kinase domain (KD), ubiquitin-like domain (ULD), scaffold/dimerization domain (SDD), and NEMO-binding domain (NBD). Crystal structure analysis showed that IKKβ is activated by trans auto-phosphorylation of Ser171 and Ser181 in the activation loop of KD through higher order oligomerization [8, 9]. IKKβ phosphorylates IкB*α* at N-terminal serines, Ser32 and Ser36, which leads to its ubiquitination and proteasomal degradation. These reactions result in nuclear translocation of NF-кB and binding of its cognate кB sites in the promoters of target genes. The NF-кB signaling pathways are associated with a vast number of human diseases, including inflammatory disorders and cancer, which suggests that IKKβ can be a potentially important therapeutic target [10].

Protein folding, maintenance of proteome integrity, and protein homeostasis critically depend on a complex network of molecular chaperones, including conserved heat shock proteins (HSPs) [11, 12]. Environmental insults perturb cellular proteome homeostasis, triggering the proteotoxic stress response, or heat shock response, that includes the transcriptional up-regulation of genes encoding HSPs. HSPs are classified into several groups based on their molecular weight. For example, Hsp60, Hsp70, and Hsp90 refer to families of HSPs on the order of 60, 70, and 90 kilodaltons in size, respectively. Hsp90, one of the most conserved HSPs, is essential not only in the protective heat shock response but also due to its chaperoning function in folding client proteins to their active conformations [13]. The Hsp90 chaperon is responsible for folding of protein kinases, and Hsp90 inhibitors such as geldanamycin (GA), stimulate kinase degradation [14]. It has been reported that Hsp90 is essential for IKKβ activity [15, 16], and Hsp90 inhibition results in IKKβ degradation [15-17]. Degradation of IKKβ is mediated by two major pathways that degrade cellular proteins in cells: the ubiquitin–proteasome system (UPS) and autophagy [17-19].

Keap1 is a ubiquitin ligase which interacts with a transcription factor Nrf2, a master regulator of the antioxidant response [20, 21]. Under quiescent conditions, Nrf2 is anchored in the cytoplasm by binding to Keap1, which in turn facilitates its proteolysis by UPS [22]. Oxidative stress and electrophiles disrupt the Keap1-Nrf2 complex, dissociating Nrf2 from Keap1 and translocating it into the nucleus where it can act as a transcription factor for series of antioxidant genes to defend against oxidative stress. Keap1 also binds to IKKβ and causes its degradation through UPS [18] or autophagy [23]. There are emerging evidence showing a functional interplay between IKKβ/NF-кB and Keap1/Nrf2 signaling pathways, which modulates inflammatory and carcinogenic processes.

Inhibition of HSPs or heat shock leads to proteotoxic stress that causes cellular protein damage or conformational changes such as protein misfolding, resulting in protein degradation. IKKβ degradation caused by Hsp90 inhibition may reflect an important adaptive cellular response which links proteotoxic stress response to inflammatory stimuli. However, the mechanism of IKKβ degradation by Hsp90 inhibition is yet be studied. In this study, we investigated the mechanism by which Hsp90 inhibition causes IKKβ degradation and found that structural destabilization of IKKβ enhances its binding to Keap1 and promotes its autophagic degradation by different mechanism from that of Nrf2.

## 2 Material and methods

### 2.1 Reagents and antibody

Anti-tubulin (H-300) and anti-HA (Y-11) rabbit antibodies were obtained from Santa Cruz Biotechnology. Anti-HA (11867423) rat monoclonal antibody and anti-Keap1 (10503-2-AP) rabbit antibody were purchased from Roche and Proteintech, respectively. Anti-IKKβ (2684), anti-phospho-IKKβ (2697) and anti-phospho-IкB*α* (2859) rabbit antibodies were procured from Cell Signaling. Anti-Flag (M2) mouse antibody was obtained from Sigma. MG132, Chloroquine, GA, 3-methyladenine(3-MA), and cycloheximide (CHX) were procured from Sigma-Aldrich. Tertiary Butylhydroquinone (tBHQ) was purchased from Tokyokasei.

### 2.2 Cell culture, plasmid transfection, and luciferase assay

Human embryonic kidney (HEK293) and mouse embryonic fibroblast (MEF) cells were cultured in Dulbecco’s modified Eagle’s medium supplemented with 10% fetal bovine serum, 2 mM L-glutamine, 100 U/ml penicillin G, and 100 μg/ml streptomycin. Keap1 and Nrf2 were amplified from a human cDNA library by polymerase chain reaction (PCR), and cDNAs were cloned in expression vectors encoding HA and Flag sequences. Expression plasmids encoding IKKβ, IKK*α*, NEMO/IKK*γ* and IкB*α* were described previously [24]. Substitution mutants were generated by PCR. Plasmids were transfected into fibroblasts, which were cultured in Opti-MEM (Invitrogen) using Lipofectamine Plus (Invitrogen), following the manufacturer’s instructions. For luciferase assay, cells were transfected with reporters encoding NF-кB binding sites (pNF-кB luciferase plasmids), or Nrf2 binding sites (pNQO1 luciferase plasmids), together with pRK-TK Renilla-luciferase control plasmids. Luciferase activity was measured using Dual-Luciferase Reporter Assay System (Promega) following the manufacturer’s instructions.

### 2.3 Cell fractionation

Cells were solubilized in buffer A consisting of 20 mM Tris-Cl (pH 7.5), 150 mM NaCl, 10 mM EGTA, 10 mM MgCl2, 60 mM β-glycerophosphate, 1 mM Na3VO4, 1 mM 4-amidino phenyl methyl sulfonyl fluoride, 50 KIU/ml aprotinin, 20 μg/ml pepstatin, 20 μg/ml leupeptin, 2 mM dithiothreitol and 1% Triton X-100. After centrifugation at 16,000 x *g* for 20 min at 4°C, the lysates were fractionated into supernatants and pellets. The pellets were then solubilized in buffer B containing 100 mM Tris-Cl, pH 6.8, 2% SDS, 5% glycerol, and 2.5% 2-mercaptoethanol. Cells solubilized in buffer B were used as total cell lysates.

### 2.4 Immunoprecipitation and immunoblotting

For the immunoprecipitation assay of transfected cells, cell supernatants were incubated with anti-Flag (M2) Sepharose (Sigma), or with an antibody combined with Protein A and Protein G Sepharose (GE Healthcare) at 4°C. For the immunoprecipitation assay of endogenous proteins, cell lysates were incubated with an antibody together with TrueBlot anti-mouse or anti-rabbit IP beads (eBioscience) and subjected to immunoblotting using TrueBlot HRP-conjugated anti-mouse or anti-rabbit IgG antibodies (eBioscience). Proteins separated by gel electrophoresis were transferred to polyvinylidene difluoride membranes (Millipore) by an electroblotting apparatus (Mighty Small Transphor; Amersham) and subjected to immunoblotting using HRP-conjugated anti-mouse or anti-rabbit IgG antibodies (GE Healthcare). Antigen-antibody complexes were detected using SuperSignal West Pico Chemiluminescence System (Pierce).

## 3 Results

### 3.1. Inhibition of Hsp90 promotes Keap1-mediated autophagic degradation of IKKβ through enhancing Keap1 binding

When cells were treated with GA, the decrease in IKKβ levels was slower in *Keap1*^*-/-*^ MEF cells than in their wild-type counterparts, suggesting that Keap1 is partially involved in GA-induced IKKβ degradation (Fig. 1A). Inhibitors of autophagy such as chloroquine and 3-MA, but not proteasome inhibitor MG132, delayed GA-induced degradation, indicating that autophagy mainly mediates GA-induced IKKβ degradation (Fig. 1B). Misfolded proteins are known to form detergent-insoluble aggresomes. GA induces translocation of transfected HA-IKKβ into detergent-insoluble fractions (Fig. 1C). IKKβ is composed of KD, ULD, SDD, and NBD, and the Keap1 binding motif which includes two essential glutamate residues (E36 and E39) locates in the N-terminal of KD (Fig. 1D). Immunoprecipitation assay revealed that GA increases binding of IKKβ to Keap1 (Fig. 1E). Glu-to-Ala substitution mutations at E36 and E39 (E36A and E39A) decreased GA-mediated enhanced IKKβ-Keap1 association (Fig. 1F). Besides, these mutations reduced the binding of co-expressed IKK*α* and NEMO/IKK*γ* to Keap1. GA-mediated degradation of IKKβ mutants (E36A and E39A) were slower than that of wild type IKKβ (wt IKKβ), suggesting that inhibition of Hsp90 promotes autophagic degradation of IKKβ mediated in part by binding to Keap1

**Fig. 1.**
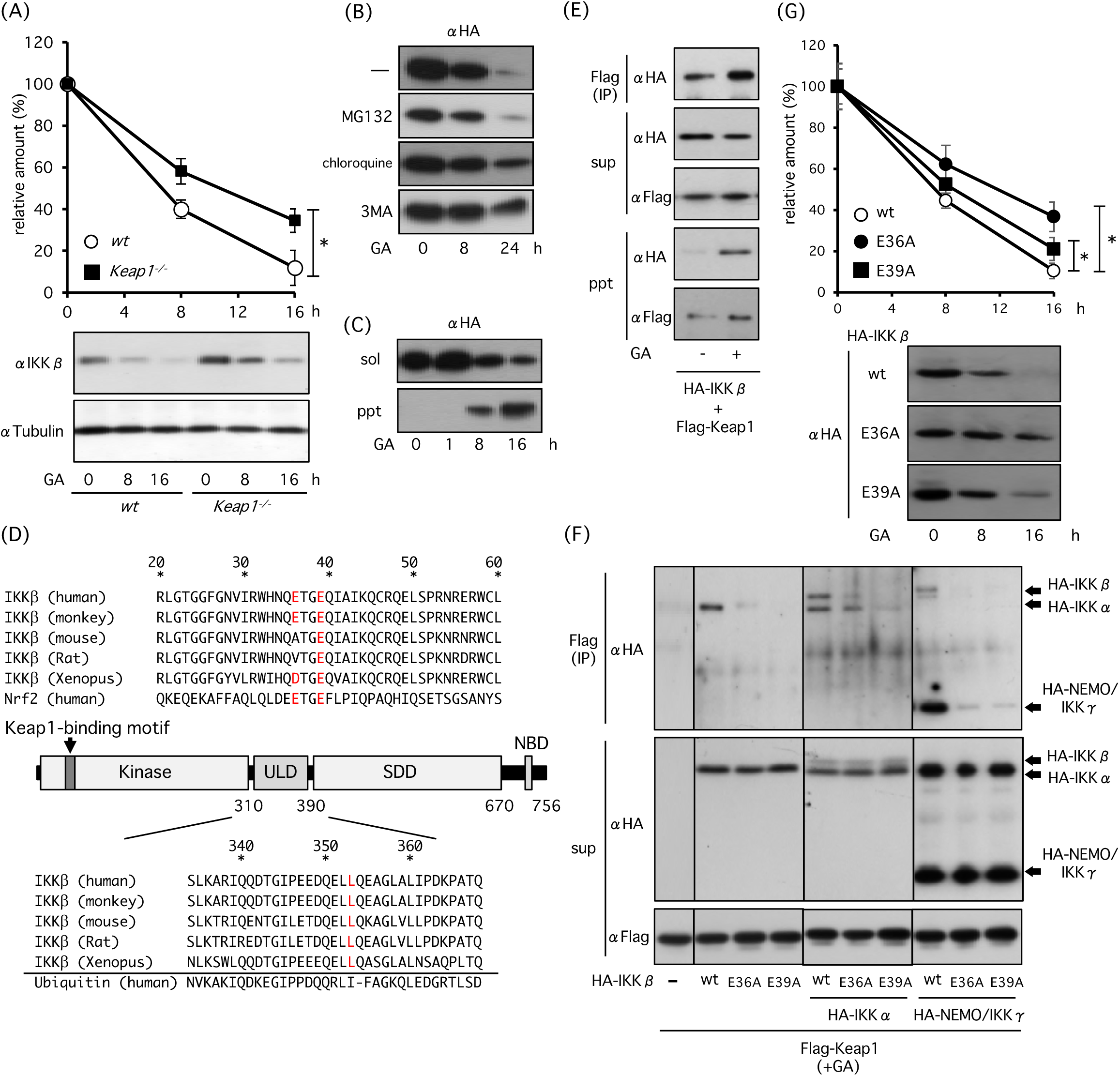
Effect of Hsp90 inhibition on IKKβdegradation. (A) Role of Keap1 in GA-induced IKKβ degradation. WT and *Keap1*^*-/-*^ MEFs were incubated with 2 µM GA, and expression levels of IKKβ were investigated. (B) Role of autophagy in GA-induced IKKβ degradation. HEK293 cells were transfected with HA-IKKβ and incubated for 18 h. Then cells were treated with GA in the presence or absence of 20 µM MG132, 20 µM chloroquine, and 4 mM 3MA. (C) Role of GA in IKKβ translocation into detergent soluble or insoluble fractions in HA-IKKβ-transfected HEK293 cells. (D) Schematic representation of IKKβ. Keap1-binding ETGE motif of IKKβ is located in the KD. (E, F) Role of E36 and E39 glutamate residues of IKKβ ETGE motif in GA-enhanced IKKβ-Keap1 association. After 18 h transfection of HEK293 cells with Flag-Keap1 and HA-IKKβ (E) or with HA-IKKβ, HA-IKK*α* and HA-NEMO/IKK*γ* (F), HEK293 cells were treated with GA for 4 h and then proteins were immunoprecipitated with anti-Flag agarose. (G) Involvement of E36 and E39 in GA-induced degradation. After 18 h transfection of HEK293 cells with HA-IKKβ mutants, cells were treated with GA. *Statistically significance at P < 0.05.

### 3.2 Keap1-mediated IKKβ degradation is not inhibited by tBHQ

Cysteine residues in Keap1 are modified by electrophiles, such as tBHQ, or reactive oxygen species, leading to Keap1 dissociation from Nrf2. As a result, Nrf2 is stabilized and it subsequently translocates to the nucleus. To elucidate whether tBHQ stabilizes IKKβ, we investigated degradation of IKKβ in the presence or absence of tBHQ, in cells treated with GA, and found that tBHQ did not prevent GA-induced IKKβ degradation (Fig. 2A). Co-transfection assay revealed that tBHQ prevented Keap1-induced Nrf2 degradation, but not IKKβ degradation (Fig. 2B). tBHQ promoted Nrf2 activity and prevented Keap1-mediated suppression of Nrf2 (Fig. 2C). However, it did not inhibit Keap1-mediated suppression of IKKβ-induced NF-кB activation (Fig. 2D). Treatment of cells with GA suppressed TNF*α*-induced NF-кB activation, and tBHQ did not prevent GA-mediated NF-кB suppression (Fig. 2E). Co-transfection with IKKβ significantly activated NF-кB, and GA suppressed IKKβ-induced NF-кB activation (Fig. 2F). GA-mediated NF-кB suppression was not prevented by tBHQ in IKKβ-transfected cells. However, GA did not suppress NF-кB-activation in cells transfected with RelA, suggesting that GA specifically suppressed IKKβ activity (Fig. 2G). Western blot analysis revealed that GA suppressed TNF-induced IKKβ phosphorylation, and tBHQ did not inhibit GA-mediated IKKβ suppression (Fig. 2H). These results indicate that Hsp90 inhibition leads to Keap1-mediated autophagic degradation that is not inhibited by tBHQ.

**Fig. 2.**
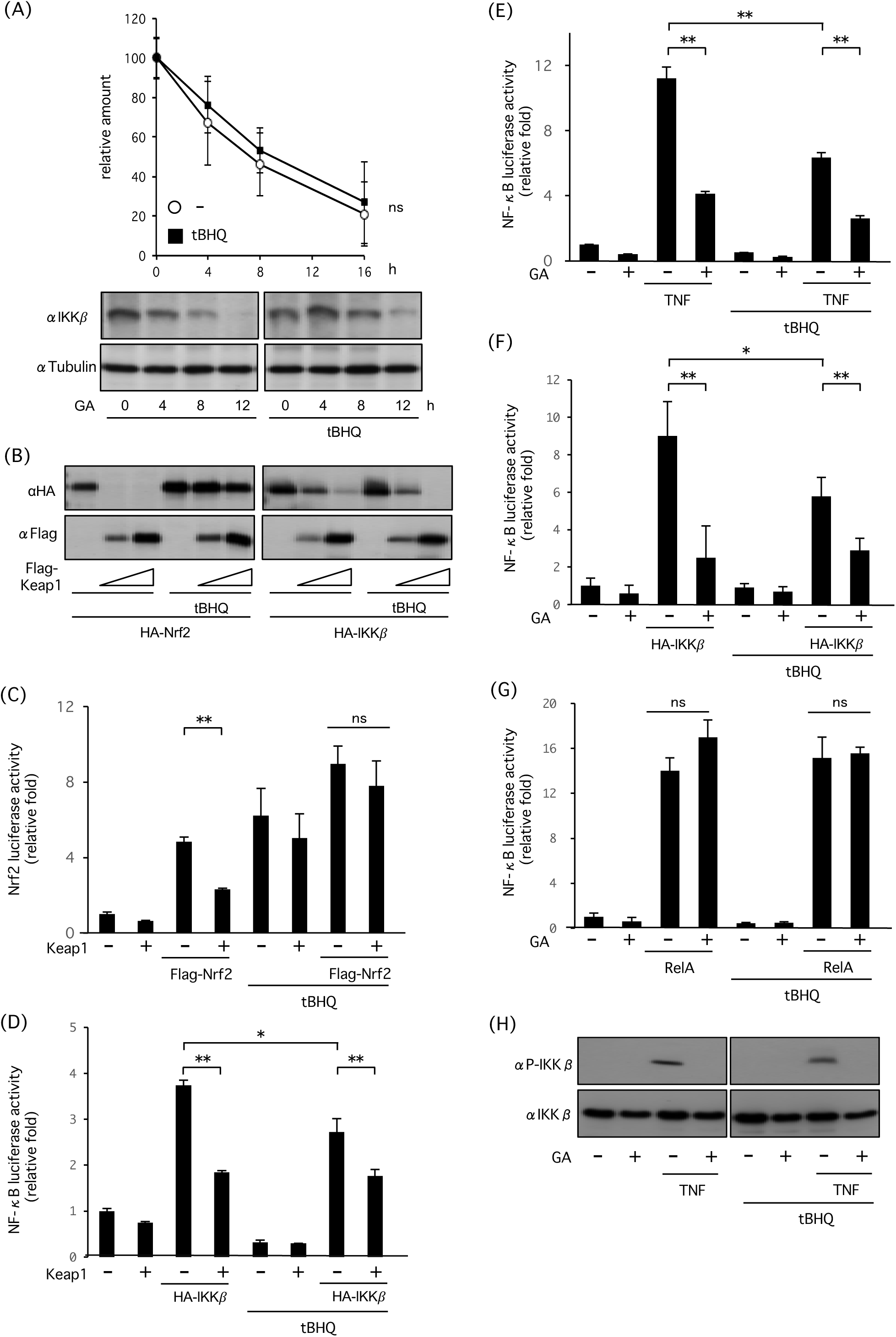
Inhibitory effect of tBHQ on Keap1-mediated IKKβ degradation. (A) Inhibitory effect of tBHQ on GA-mediated IKKβ degradation. Cells were treated with GA in the presence or absence of 50 µM tBHQ. (B) Inhibitory effect of tBHQ on Keap1-mediated Nrf2 or/and IKKβ degradation. HEK293 cells were transfected with HA-IKKβ, HA-Nrf2 and Flag-Keap1, and incubated with or without tBHQ for 18 h. (C) Nrf2 luciferase activity of HEK293 cells transfected with Nrf2-reporter together with Flag-Keap1 and HA-Nrf2 in the presence or absence of tBHQ. (D) NF-κB luciferase activity of HEK293 cells transfected with NF-κB-reporter together with Flag-Keap1 and HA-IKKβ in the presence or absence of tBHQ. Both luciferase activities were measured 18 h post-transfection. (E) NF-κB luciferase activity of HEK293 cells transfected with NF-κB reporter, and incubated with or without GA, tBHQ, and 50 ng/ml TNFα (as an inflammation stimulator) for 18 h. (F, G) NF-κB luciferase activity of HEK293 cells transfected with NF-κB reporter together with or without HA-IKKβ(F) or Flag-RelA (G), incubated in presence or absence of GA and tBHQ for 18 h. (H) Inhibitory effect of tBHQ on GA-mediated IKKβ inactivation and degradation. HEK293 cells were treated with GA for 6 h and then stimulated with TNFα for 20 min. * P < 0.05, ** P < 0.01.

### 3.3 L353A mutation destabilizes IKKβ and leads its autophagic degradation

Structural destabilized mutant of IKKβ was studied to elucidate whether structural destabilization per se induces autophagic degradation. The ULD interacts with KD and SDD in a highly organized manner that is required for a critical interaction with IкB*α* and kinase activity [8]. We generated several Ala substitution mutants in the ULD and found that Leu-to-Ala substitution mutation at Leu353 abolishes IKKβ-induced NF-кB activation (Fig. 3A), which was consistent with previous reports [8, 25]. Moreover, autophosphorylation was reduced in L353A mutant (Fig. 3B). Co-transfection of HEK293 cells with HA-IKKβ mutants and ubiquitination-resistant HA-IкB*α*RR mutant revealed that L353A mutant hardly phosphorylate IкB*α* (Fig. 3C). L353A mutant was localized to the detergent-insoluble fraction (Fig. 3D). L353A mutant degraded more rapidly than wt IKKβ, and the rate of disappearance was faster in the detergent-insoluble fraction than in the detergent-soluble fraction (Fig. 3E). Degradation of L353A mutant was partially prevented by chloroquine but not by MG132, suggesting that autophagy mainly mediates degradation of structural destabilized L353A mutant (Fig. 3F).

**Fig. 3.**
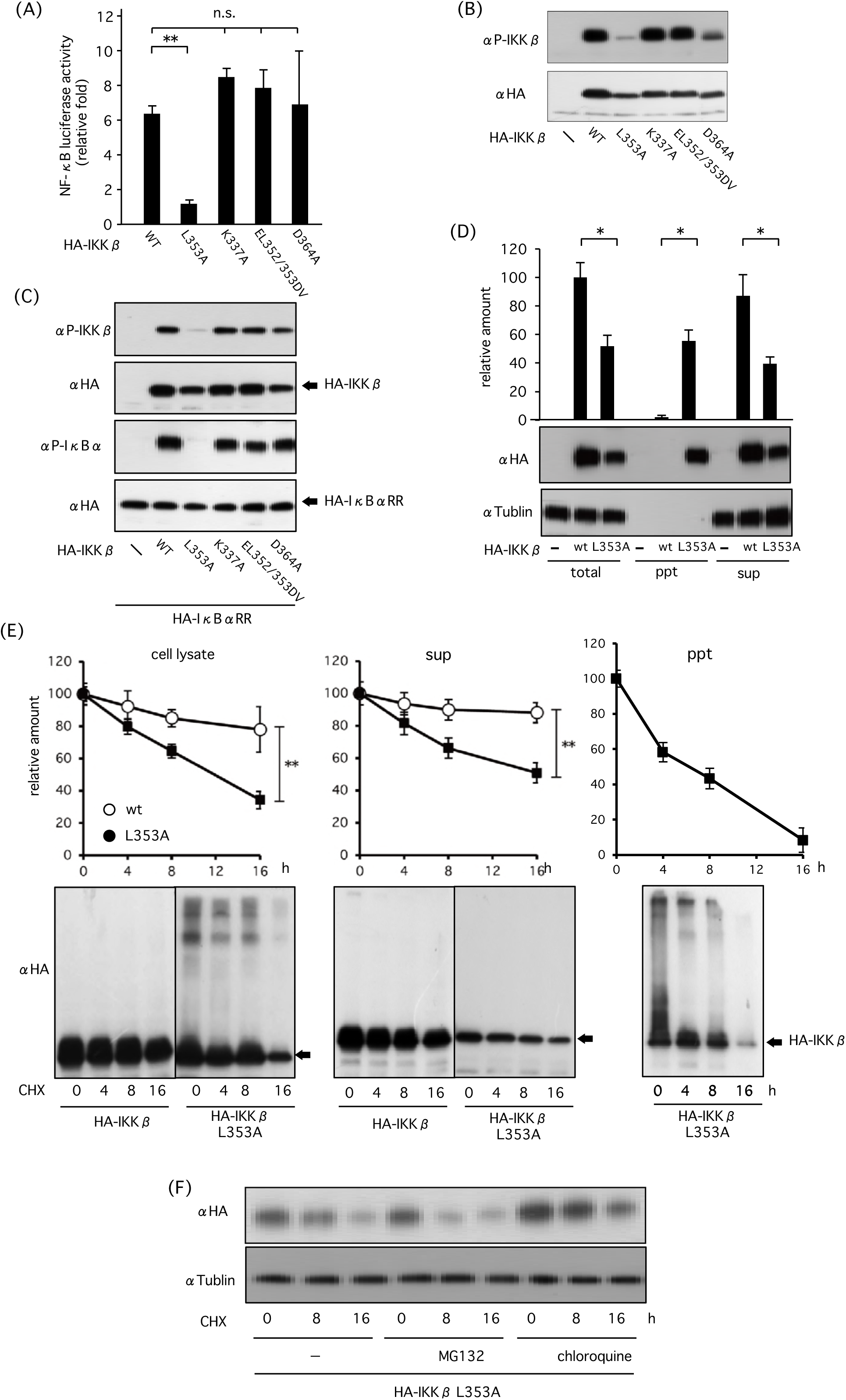
Effect of L353A mutation in the ULD of IKKβ on structural stability or degradation of IKKβ. (A) Effect of different HA-IKKβ mutations on IKKβ-induced NF-κB activation in terms of NF-κB luciferase activity. HEK293 cells were transfected with NF-κB reporter and HA-IKKβ mutants. After 18 h, luciferase activities were measured. (B) Effect of HA-IKKβ mutants on IKKβ autophosphorylation. HEK293 cells were transfected with HA-IKKβ mutants and phosphorylation of IKKβ was detected by western blot analysis. (C) Effect of HA-IKKβ on IKKβ-induced IκBα phosphorylation. HEK293 cells were transfected with HA-IKKβ mutants together with HA-IκBαRR, and phosphorylation of IκBα was detected by western blot analysis. (D) Role of L353A mutation in HA-IKKβ translocation into detergent insoluble or insoluble fractions of HEK293 cells. Cells were transfected with wild type or mutated HA-IKKβ(L353A), and expression levels were investigated in total cell lysates, detergent soluble and insoluble fractions. (E) Effect of L353A mutation on IKKβ degradation in different fractions. HEK293 cells were transfected with wild type or mutated HA-KKβ(L353A) mutant. After incubation with 10 µg/ml CHX, expression levels were investigated in total cell lysates, detergent soluble and insoluble fractions. (F) Autophagy mediates degradation of L353A mutant. HEK293 cells transfected with HA-IKKβ or HA- KKβ(L353A) mutants were incubated with CHX in the presence or absence of MG132 and chloroquine. * P < 0.05, ** P < 0.01.

### 3.4. L353A mutation increases Keap1 binding

To elucidate whether Keap1 is involved in degradation of L353A mutant, we investigated the interaction of Keap1 to L353A mutant. Immunoprecipitation assay of transfected cells revealed that binding of Keap1 to IKKβ is increased in L353A mutant, whereas binding of IKK*α*, IKKβ and NEMO/IKK*γ* to IKKβ were not changed (Fig. 4A). Additionally, the increased interaction of endogenous Keap1 to L353A mutant was detected in cells (Fig. 4B). The reduction in the amount of L353A mutant was slower in *Keap1*^*-/-*^ than in wt MEFs under CHX treatment, suggesting that Keap1 is involved in L353A degradation (Fig. 4C). Co-transfection assay revealed that Keap1 degraded L353A mutant more effectively than wt IKKβ (Fig. 4D). We generated expression plasmids encoding double mutants of L353A/E36A and L353A/E39A. L353A/E36A was expressed effectively, whereas L353A/E39A was hardly expressed in cells. Then we compared degradation ratio of L353A/E36A to L353A and found that L353A/E36A is more stable than L353A (Fig. 4E). These results indicate that structural destabilization of IKKβ increases its interaction to Keap1, leading to autophagic degradation.

**Fig. 4.**
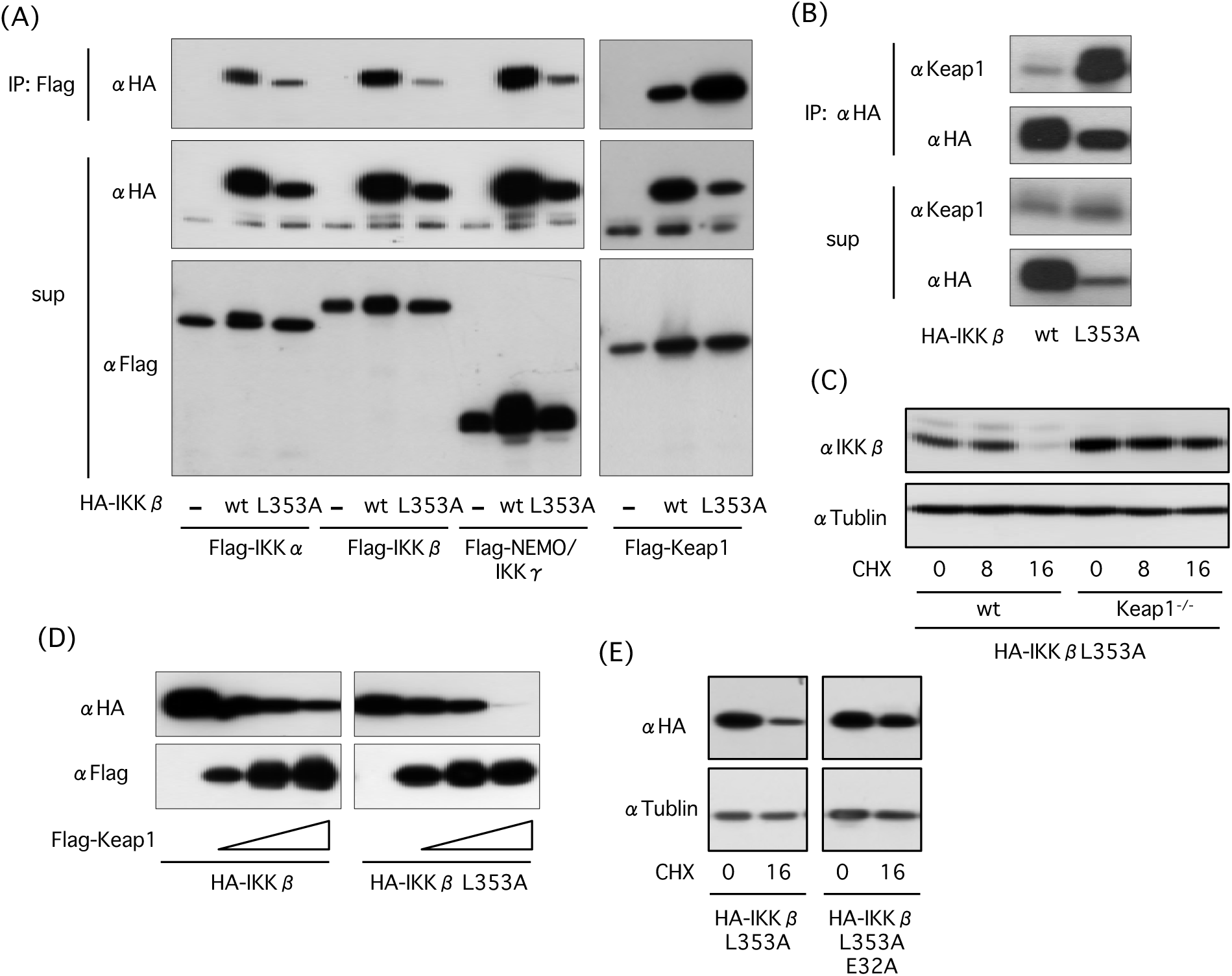
Effect of L353A mutation on association of IKKβ with Keap1 and autophagic degradation. (A) Effect of L353A mutation on IKKβ-Keap1 binding. Cells were transfected with wild type or mutated HA-IKKβ(L353A) together with Flag-IKKα, Flag-IKKβ, Flag-NEMO/IKKγ, and Flag-Keap1. After 18 h, cell lysates were immunoprecipitated with an anti-Flag agarose beads, and association of IKKβ with IKKα, IKKβ, NEMO/IKKγ, and Keap1 was investigated. (B) Effect of L353A mutation on association of IKKβ with endogenous Keap1. HEK293 cells transfected with HA-IKKβ or HA-IKKβ(L353A) were immunoprecipitated with an anti-HA antibody. (C) Role of Keap1 in IKKβ(L353A) degradation. After transfection of WT and *Keap1*^*-/-*^ MEF cells with mutated HA-IKKβ(L353A), cells were treated with CHX and degradation of HA- KKβ(L353A) was investigated by western blot analysis. (D) Effect of L353A mutation on Keap1-mediated IKKβ degradation. HEK293 cells were transfected with HA-IKKβ or HA-IKKβ(L353A) together with Flag-Keap1, and expression levels were investigated in cell lysates by western blot analysis. (E) Involvement of Keap1-binding motif in degradation of L353A mutant. After 18 h transfection with HA-IKKβ(L353A) and double mutant HA-IKKβ (L353A/E36A), HEK293 cells were incubated with CHX HA expression was detected by western blot analysis.

## 4 Discussion

The IKKβ/NF-кB and Keap1/Nrf2 signaling pathways play an essential role in cellular signaling in response to inflammatory stimuli and oxidative stress. It has been reported that activation of NF-кB antagonizes Nrf2-mediated transcription by either inhibition of Nrf2-DNA binding [26] or depriving CREB binding protein (CBP) from Nrf2 and facilitating the interaction of recruited histone deacetylase 3 (HDAC3) at the Nrf2-DNA binding site [27]. Nrf2-deficient MEFs show greater activation of NF-кB in response to lipopolysaccharide (LPS) stimulus [28], and Nrf2 suppresses NF-кB-induced transcriptional upregulation of proinflammatory cytokines, including IL-6 and IL-1β in macrophages [29]. Indeed, NF-кB activation is suppressed in tBHQ-treated cells in which Nrf2 is activated (Fig. 2D, E, and F). Thus, cellular responses to inflammatory stimuli and oxidative stress is regulated by crosstalk between the IKKβ/NF-кB signaling and the Keap1/Nrf2 signaling pathways.

Although Keap1 is involved in structurally destabilized IKKβ degradation, the contribution of Keap1 to degradation is partial; thus, other factors would be involved in IKKβ degradation as well. Recent study has revealed that Drosophila Kenny, the homolog of mammalian NEMO/IKK*γ*, is a selective autophagy receptor that mediates the degradation of the IKK complex [30]. Drosophila NEMO/IKK*γ* has an LC3-interacting region (LIR) or Atg8-interacting motif (AIM) in the N-terminal region which recruits the IKK complex to the autophagosome. This selective autophagic degradation of the IKK complex prevents constitutive activation of the immune deficiency pathway in response to commensal microbiota in Drosophila [30]. Remarkably, human NEMO/IKK*γ* lacks functional LIR/AIM and does not interact with mammalian Atg8-family proteins. Instead, NEMO/IKKγ in mammals has been shown to be ubiquitinated, and therefore, could also interact with selective autophagy receptors via ubiquitin tags [31, 32]. Thus, several mechanisms would be involved in IKKβ degradation in cells under proteotoxic stress.

Cells are challenged by various extrinsic and intrinsic insults which provoke proteotoxic stress under conditions of injury, inflammation, and oxidative stress. Proteome homeostasis is essential to maintain cellular fitness and its disturbance is associated with a broad range of human diseases. To protect cells from proteotoxic stress, cells mobilize heat shock proteins including Hsp90 that unfold proteins and proteolysis systems that remove damaged proteins. In this study, it was found that Hsp90 inhibition, which causes proteotoxic stress, enhances interaction of IKKβ with Keap1. These findings also suggest that cells respond to proteotoxic stress by regulating IKKβ/NF-кB and Keap1/Nrf2 signaling pathways. The Keap1/Nrf2 and IKKβ/NF-кB signaling pathways have interesting features concerning the proteolytic mechanisms such as autophagy and UPS. Keap1 suppresses Nrf2 activation by UPS, whereas it is degraded by autophagy in response to oxidative stress [33, 34]. IKKβ induces NF-кB activation through degradation of IкB*α* by UPS, whereas it is degraded by autophagy in response to proteotoxic stress. Furthermore, IKKβ contributes to the induction of autophagy in an NF-кB-independent manner [35], and Keap1 interacts with p62, a selective autophagy receptor, which binds to Atg8/LC3 protein, resulting in autophagy [36, 37]. There is a plausibility that oxidative stress and proteotoxic stress induce selective degradation of Keap1 and IKKβ by autophagy and provoke specific cellular responses.

## 5. Acknowledgments

This work was supported, in whole or in part, by Grants-in-Aid from the Ministry of Education, Culture, Sports, Science, and Technology of Japan (#15H01406, # 15H01382, and #23370062). The content is solely the responsibility of the authors and does not necessarily represent the official views of the Ministry of Education, Culture, Sports, Science, and Technology of Japan.

